# Analysis of infection biomarkers within a Bayesian framework reveals their role in pneumococcal pneumonia diagnosis in HIV patients

**DOI:** 10.1101/070144

**Authors:** Austin G. Meyer

**Affiliations:** Center for Computational Biology and Bioinformatics, The University of Texas at Austin, Austin, TX, USA; School of Medicine, Texas Tech University Health Sciences Center, Lubbock, TX, USA

## Abstract

**Background:** HIV patients are more likely to contract bacterial pneumonia and more likely to die from the infection. Unfortunately, there are few tests to quickly diagnosis the etiology of these dangerous infections. Several biomarkers may be useful for diagnosing the most common pneumonia-causing organism, *S. pneumoniae,* but studies utilizing the standard statistical approach provide little concrete guidance for the HIV-infected population.

**Methodology and Findings:** Using a Bayesian approach, I analyze data from a cohort of 280 HIV patients with x-ray confirmed community acquired pneumonia. First, I use a variety of techniques to establish predictor significance and to identify their optimal cutoffs. Next, in lieu of cutoffs, I find the continuous and combined likelihood ratios for every value of each biomarker, and I compute the associated posttest probabilities. As expected, I find the three biomarkers with good clinical yield and a statistically significant association with *S. pneumoniae* are C-reactive protein (CRP), procalcitonin (PCT), and *lytA* gene PCR (lytA). Based on Bayesian clinical yield, optimal cutoffs are largely equivocal. The optimal dichotomous cutoff for CRP is essentially any value between 10 *mg/dL* and 30 *mg/dL* (△*p_Posttest_* ≈ 0.49). The optimal cutoff for PCT is any value between 2 *ng/mL* and 40 *ng/mL* (△*p_posttest_* ≈ 0.35). The optimal cutoff for lytA is any value less than 6 *log*_10_ *copies/mL* (△*p_posttest_* ≈ 0.45). Further, I find that continuous likelihood ratios provide much more accurate posttest probabilities than dichotomous cutoffs. For example, starting with the empirical pretest probability, a lytA approaching 0 *copies/mL* lowers the probability of S. *pneumoniae* infection to less than 15%, while a result of 10 *copies/mL* raises the probability to greater than 65%. However, a lytA value just above or below the suggested cutoff of 8000 *copies/mL* or my new optimal cutoff of 30,000 *copies/mL* leaves the posttest probability of infection essentially unchanged from the pretest probability.

**Conclusion:** CRP, PCT, and lytA all provide significant value in diagnosing the etiology of pneumonia in HIV patients. The optimal dichotomous cutoffs for lytA, CRP, and PCT need to be adjusted for pneumococcal diagnosis in this population. However, continuous and combined likelihood ratios avoid discarding valuable quantitative information, and a combined likelihood ratio can be easily computed without the need for prior logistic regression. Importantly, there is significant overlap between these biomarkers such that only one of the three biomarkers at a time should be used to update clinical probabilities. Thus, it is ill-advised to combine the likelihood ratios of different biomarkers to produce a posttest probability. Finally, I provide a simple web application to quantitatively calculate the posttest probability of *S. pneumoniae* infection in HIV patients with x-ray confirmed pneumonia: http://meyerapps.org/pneumococcal_etiology_hiv.

## Introduction

Respiratory infections are among the most common causes of death and disease worldwide, and regions with a high incidence of HIV are the hardest hit. HIV infected individuals are at least twenty five times more likely to develop a bacterial pneumonia compared to their uninfected peers [1, 2]. Approximately sixty percent of acute lower respiratory tract infections in HIV patients have a bacterial etiology, and seventy percent of those infections are caused by *Streptococcus pneumoniae* [3]. Despite the etiologic concordance between HIV-infected and uninfected populations, the morbidity and mortality from invasive *S. pneumoniae* infection is much higher in HIV patients. Some demographic subgroups are thirty to one hundred times more likely to develop disseminated pneumococcal disease following pneumococcal pulmonary infections [2,4]. Nevertheless, there are many other respiratory pathogens that frequently infect HIV patients [2,3]. The most common bacteria include *Haemophilus influenzae, Staphylococcus aureus*, and *Legionella pneumophila* each with their own local variation in empiric treatment. Mycobacteria species including *M. tuberculosis* account for nearly twenty percent of pulmonary infections in HIV patients while viruses, fungi, and parasites make up the remaining ten percent of infections [3]. In every case, it is important to rapidly diagnose the infecting organism both for treatment success and for antimicrobial stewardship.

There are many biomarkers with clinical potential for rapidly evaluating pneumonia etiology [5,6]. To identify infections and hint at an organism, physicians follow core body temperature, white blood cell count and the erythrocyte sedimentation rate [7]. Unfortunately, these markers are considered relatively non-specific for particular organisms. More recently, there are a number of biomarkers referred to collectively as acute phase reactants that have been validated for a wide range of clinical applications [7]. Among the most common is C-reactive protein (CRP), a protein that is synthesized in the liver, and was originally identified as a serological fraction in patients with *S. pneumoniae*-infected patients [7–10]. Despite its identification in pneumococcal infections, CRP became known as a non-specific inflammatory marker with application in a wide range of disciplines from rheumatology to cardiology [7]. For respiratory infections, studies show that CRP levels tend to be higher in *S. pneumoniae* and *L. pneumophila* pulmonary infections compared to infections by atypical organisms [11]. Furthermore, several studies suggest that a high CRP can be used to determine disease severity for bacterial infections [8,12]. Despite some success, CRP retains its stigma as a non-specific marker of inflammatory disease. A different acute phase biomarker, procalcitonin (PCT), appears to be more valuable then CRP for the evaluation of acute respiratory infections [13–15]. Several randomized control trials show that PCT can be used both to track disease status and to guide antibiotic administration [16]. In contrast to CRP, studies generally demonstrate that PCT is relatively specific for bacterial infections. The evidence for PCT is so overwhelming that the federal Agency for Healthcare Research and Quality recommends its use for initiating and discontinuing antibiotics in patients with generic respiratory infections [17]. However, as of yet, there is relatively little evidence for the use of PCT in several important populations including HIV patients and patients with cystic fibrosis [17]. In the last five years, a real time polymerase chain reaction (rtPCR) test for the pneumococcal gene *lytA* (lytA) in nasopharyngeal swabs has proven to be sensitive and specific for the colonization of *S. pneumoniae* [18–20]. Moreover, tests for nasopharyngeal lytA are significantly higher in HIV patients with pneumococcal pneumonia than in asymptomatic HIV-infected controls [21,22]. In addition, analyses of data from HIV-uninfected patients show that lytA densities of greater than 8, 000 *copies/mL* may be a useful diagnostic indicator for *S. pneumonia* pulmonary infections [23].

Despite some connection between biomarkers and pneumonia etiology, in the absence of specific randomized control trials (RCT) for every possible population and use case, it is difficult to translate most of these tests into actionable guidance. Even when there are good RCTs, it is difficult for clinicians to know how to extrapolate study results to their particular patient population. Moreover, in the vast majority of clinical studies, investigators prefer to reduce continuous quantitative tests to just two categories via a dichotomous cutoff (i.e. a test cutoff where test results are either greater than or less than the threshold value) [24,25]. That choice largely stems from the dominant approach to statistical analyses called frequentist statistics, which focuses on determining whether samples are drawn from the same or different underlying distributions. For example, in the initial case of evaluating biomarkers, investigators enroll a large number of patients with a disease (based on a gold standard confirmatory test) and a similarly large number of asymptomatic controls. Each patient is tested for the biomarker. Then, investigators use a statistical test to determine whether patients and controls are drawn from the same underlying biomarker distribution; assuming they are from the same distribution, researchers compute the probability (the p—value) of the identified level of extremeness by chance alone. Once the biomarker test is statistically significant by p—value, they use other metrics to identify the clinical cutoff that maximizes the true positive rate and simultaneously minimizes the false positive rate [24–26]. Thus, the practice of frequentist inference revolves around quantifying the probability of incorrectly classifying a person given their test result.

For clinical applications, there are at least three major problems with the frequentist approach. First, for most people frequentist probabilities are not a natural way to reason. For individual patients, a clinician wants to know the probability that a patient has a particular disease; at best, a standard frequentist analysis only computes the probability that the patient is misclassified as diseased. Second, since frequentist statistics generally find the probability of sample extremeness, reasoning with frequentist statistics is most natural and convenient with dichotomous cutoffs (e.g. disease and not disease) [26]. Therefore, the approach implies that clinicians and researchers should throw out possibly informative quantitative differences between patients who may fall on the same side of the dichotomous threshold. There is no clear reason to prefer eliminating such a distinction [27–30]; if anything, there are good reasons to prefer retaining the additional information in quantitative biomarkers. Third, frequentist inference eliminates the possibility of easily defining a prior probability [31]. Certainly a 60 year-old HIV patient with a CD4 count of 50 *cells/mL*, with a 100 pack-year smoking history, and a viral load of 100, 000 *copies/mL* on no prophylaxis is not at the same risk for pneumonia as a 30 year-old with CD4 count of 700 *cells/mL*, with no smoking history, and an undetectable viral load. Therefore, even though it is comparatively easy to calculate empirical prior probabilities for many diseases, incorporating that information is essentially impossible.

By contrast, a Bayesian statistical approach easily answers all of these challenges. First, Bayesian probabilities are the natural way that humans reason. Clinicians always implicitly begin with a prior probability of disease [31]. In fact, the practice of building and ranking a differential diagnosis is founded on the Bayesian principle of prior probability. Next, clinicians perform a test and subsequently update their degree of belief that a patient has a particular disease [27,28,31]. Mathematically, a likelihood ratio is the most convenient means to update the probability of disease [31]. Although most Bayesian analyses of biomarker tests fall into the same dichotomous trap as their frequentist contemporaries, that is certainly not necessary or particularly helpful. One can calculate likelihood ratios for every possible test result, and those continuous likelihood ratios can be used to update the pretest probability of disease for individual patients [26,27,29]. Then, clinicians are left with a simple, interpretable probability that represents the probability of disease.

In this study, I find that CRP, PCT, and lytA can all independently aide in the diagnosis of pneumococcal pneumonia, but they cannot be combined to produce improved results. In addition, the current cut points for each test do not optimize their discriminatory power. According to the best traditional method and a more modern Bayesian approach, the cutoffs should be increased for etiology determination. Nevertheless, optimizing the cutoff is probably not ideal for clinical utility. Instead, combined and continuous likelihood ratios can be easily computed and retain all of the quantitative information inherent in the tests. I find that retaining the quantitative information improves the accuracy of posttest probabilities relative to setting a single cutoff. Last, I provide a web application that allows clinicians to quickly input test results and compute a posttest probability of *S. pneumoniae* respiratory infection. Although, the web application serves a single defined purposed, my hope is that a similar approach can be easily incorporated into lab result reporting systems within existing electronic medical record applications.

## Methods

The data used in this study was collected and analyzed previously in a separate manner [22]. I downloaded the data from Data Dryad [32]. The data included a total of 280 HIV infected patients. Patients were admitted to the study after having a confirmatory chest x-ray for community acquired pneumonia. There were many available predictors in the dataset including age, Bartlett score, CD4 count, Bactrim prophylaxis status (either taking or not taking), HAART therapy status (either taking or not taking), CURB65 score, proANP, proADM, copeptin, procalcitonin, C-reactive protein, and bacteremia. In addition, there were three response variables including bacteremia (either bacteremic or not bacteremic), pneumococcal etiology by standard criteria (either confirmed or ruled out), and pneumococcal etiology by expanded criteria. Pneumococcal etiology was established if a sputum culture, Gram’s stain, urinary pneumococcal antigen, or blood culture revealed pneumococci. The dataset included an additional criteria to establish an expanded definition of pneuomococcal etiology; that criteria was a threshold of 8 × 10^3^ *copies/mL.* I chose to omit that criteria to avoid internal circularity in interpreting the results. Here, I used only pneumococcal diagnosis as the response variable.

All analyses were performed in the R statistical programming language [33]. I used several libraries to import, split, combine, filter, and reshape the data [34–37]. I used the plotting library ggplot2 to generate all of the figures in this manuscript [38]. I used the supplementary functions provided by cowplot to make final, publication-ready figures [39]. Everything not provided by these packages, I custom coded. I made available all of my code, figures, and data on Github: https://github.com/ausmeyer/pneumococcal_ colonization_analysis_redo.

I performed univariate logistic regression with every available predictor using the standard etiology criteria as the response (some results are only on Github). Then, I used age, CRP, PCT, and lytA in all pairwise combinations as predictors in a multivariate logistic model. Age was not a significant predictor, and I therefore eliminated it from all subsequent analyses. Statistical significance in multivariate regression was established with the *p*—value of each predictor variable. Next, I computed the sensitivity and specificity for every predictor value in the dataset. The sensitivity and specificity were computed as

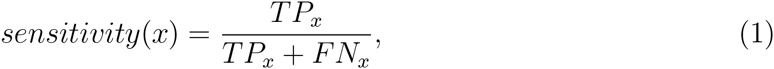

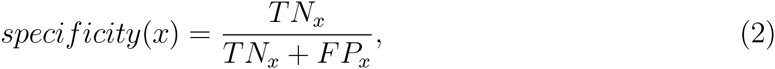

where *x* represents each biomarker value. Thus, *TP_x_* is the number of true positives at *x*, *FP_x_* is the number of false positives at *x*, *TN_x_* is the number of true negatives at *x*, and *FN_x_* is the number of false negatives at *x*. I calculated the Youden index [24,25] empirically at each biomarker value as

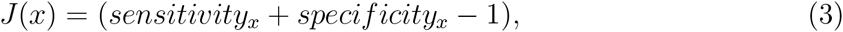

where J is the Youden J index. For the standard method, I found the ideal cutoff value for each biomarker as the biomarker value, x, that maximized the Youden *J*(x).

The transition to Bayesian statistics required computing several additional quantities [31]. The easiest and most extensible approach for clinical decision-making involves the use of likelihood ratios to update the probability of disease after administering a clinical test [31]. In an effort to compute likelihood ratios over the range of data with relatively smooth behavior, I narrowed the upper limit of CRP to 35 *mg/dL.* This design decision had no effect on the analysis other than to eliminate inappropriate edge effects in the data that resulted from the relatively small boundary sample size.

I started by calculating the empirical pretest probability in the dataset as the prevalence of pneumococcal disease. Then, the LR+ and *LR–* were calculated with a simple ratio of sensitivity and specificity relations:

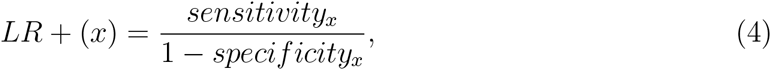

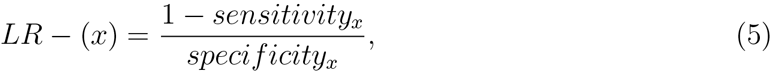

where *x* again is every value of each biomarker. The associated confidence interval was calculated in accordance with [40] as

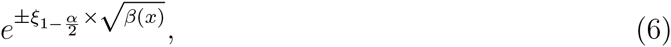

where 1 — *α* is the standard confidence level and

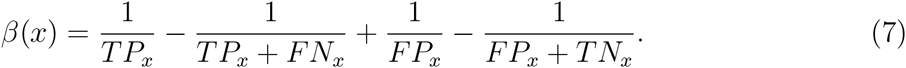

At first glance, this notation may seem overly formal. However, the 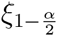 is simply the standard quantile of the normal distribution on the desired confidence interval. If that is unclear, the code is publicly available in the Github repository. Starting with the pretest probability, I found the posttest positive and negative probabilities at each possible biomarker cutoff. That required first calculating the pretest odds from the pretest probability. Then, I updated the odds by multiplying the pretest odds by the relevant likelihood ratio (either *LR*+ or *LR*–). The odds were converted back to probabilities to produce two posttest probability values at each cutoff. To find the optimal cutoff, I took the difference of the two posttest probabilties at each point with

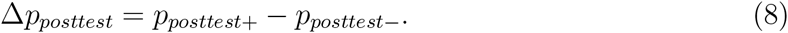

Finally, the optimal cutoff value of a biomarker is simply the test value that maximizes the difference in the two posttest probabilities.

Next, I proceeded to calculate the continuous likelihood ratio by two different means. In the first, I used the methodology of Simel et al. [26]. This method requires first fitting a logistic model [27], then computing the likelihood ratio using the logistic parameters. Thus, I used the logistic probability function,

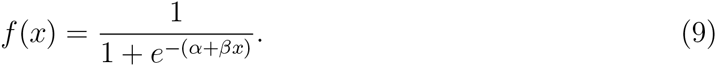

Then, with the values from the logistic fit (as in Simel et al.), I calculated the continuous likelihood ratio for each value of the biomarkers with

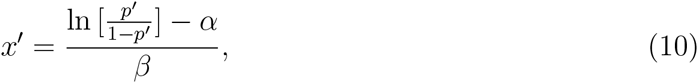

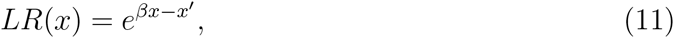

where *p′* is the pretest probability of the relevant condition. In this case, I used the empirical pretest probability. In addition, to ensure the appropriate behavior of the continuous likelihood ratio function, I computed the continuous likelihood ratio for a range of pretest probabilities from 0.05 to 0.95 in increments of 0.05 (result not shown).

Finally, owing to the multiplicative nature of likelihood ratios, I computed point estimates of the combined likelihood ratio. There could be some debate about the best way to accomplish the calculation empirically. For example, one could calculate the point estimate on an interval by multiplying the *LR*+ of a biomarker cutoff value *n* by the *LR*— for the cutoff point at *n* +1. In the end, I chose to compute the empirical likelihood ratio point estimates by multiplying the *LR*+ and *LR*– at a single cutoff value *n*. After some simplification, this equation takes the form

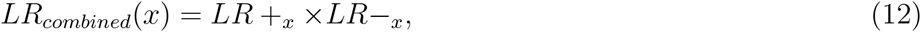

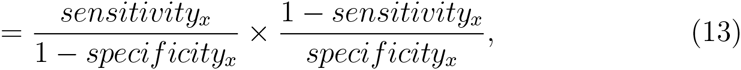

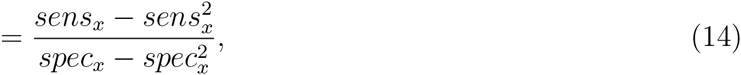

which could be further simplified into a quadratic form utilizing only *TP_x_, FP_x_, TN_x_*, and *FN_x_.* However, that simplification adds little clarity. I then repeated that calculation at every biomarker value, *x*, in the dataset. Then, in the same manner as above, I used the continuous and empirically combined likelihood ratios to update the empirical pretest probability to obtain the plotted posttest probabilities.

An accompanying website to compute the posttest probability of pneumococcal infection is available: http://meyerapps.org/pneumococcal_etiology_hiv. Code and data for the web application are freely available on Github: https://github.com/ausmeyer/pneumococcal_etiology_hiv. I built the website using the shiny server framework and the plotly interactive plotting library [41,42].

## Results

### Biomarkers can help to identify *S. pneumoniae* etiology

Univariate logistic regression revealed that only three of the predictive biomarkers had a strong connection with pneumococcal etiology (Fig. 1). These three were C-reactive protein (*n* = 41; *p_intercept_* = 1.71 × 10^−3^; *p_predictor_* = 2.56 × 10–^3^), procalcitonin (*n* = 233, *p_intercept_* = 4.46 × 10; *p_predictor_* = 4.38 × 10) > and lytA (*n* = 264, *p_intercept_*= 2.94 × 10^−14^ *p_predictor_* = 9.45 × 10^−14^). However, every pairwise combination of predictors in a multivariate logistic regression model failed to produce statistical significance in the additional predictor. Therefore, there is substantial overlap among the three predictors. From a clinical perspective, this means that only one predictor should be used to update the posttest probability of pneumococcal infection. Also, from logistic regression alone, there is no reason to prefer one of the three markers over any other.

**Figure 1:**
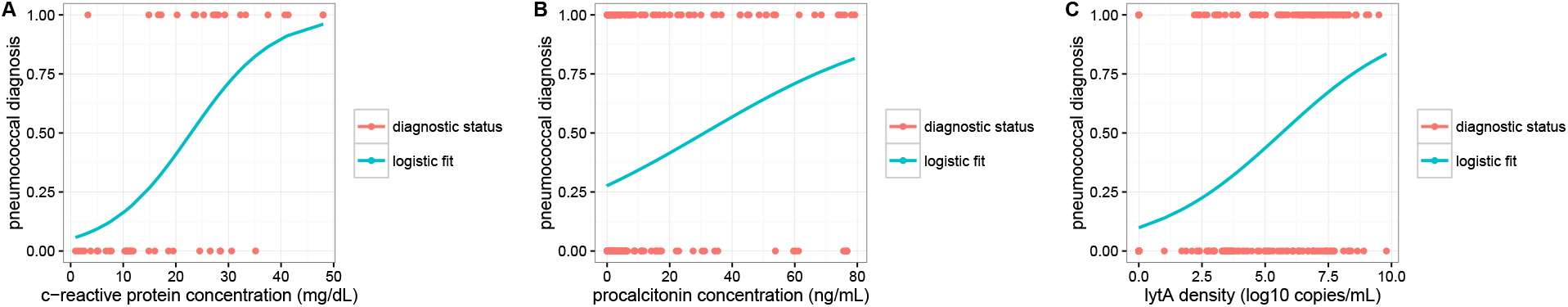
Logistic fit of real pneumococcal diagnostic status. The red points show the actual pneumococcal status of HIV patients in this study. Patients with pneuomococcal etiology are coded as 1 and those without pneumococcal etiology are coded as a 0. In A, I show the fit for C-reactive protein. In B, I show the fit for procalcitonin. In C, I show the fit for lytA.

In the interest of statistical simplicity, most modern medical tests utilize a dichotomous cutoff (a single line that divides positive tests from negative tests), even for tests that are clearly non-binary in nature. To help identify the ideal cutoff, I computed the sensitivity and specificity for each available test value in the dataset (Fig. 2). For CRP, the established “normal” range is generally less than 3 *mg/dL* or less than 10 *mg/dL* for some high sensitivity tests. By contrast, I found an equivalence point between 20 *mg/dL* and 22.5 *mg/dL* where the sensitivity and specificity were both above 75%. In the CRP “normal” range, the specificity to diagnose pneumococcal pneumonia approached zero. Similarly for procalcitonin, there was equivalence near 2 *ng/mL* where both sensitivity and specificity were approximately 75%, which is far above the normal cutoff of 0.5 *ng/mL.* For lytA, I again found an equivalence point at approximately 4.5 *log*_10_ *copies/mL* with a similar sensitivity and specificity. Thus, plotting sensitivity and specificity suggested the same conclusion as that from logistic regression; each of the tests provided similar value in diagnosing pneumococcal pneumonia. However, a different statistic is required to identify the ideal cutoff.

**Figure 2:**
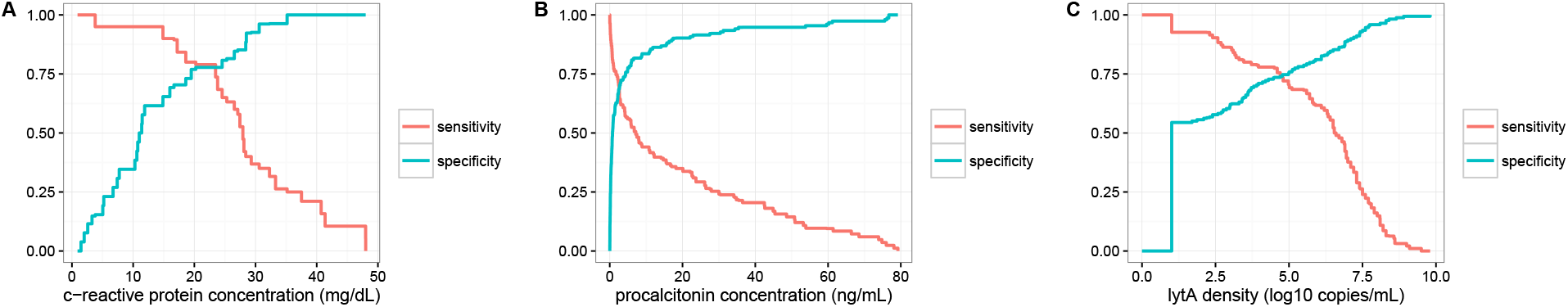
Sensitivity and specificity plots at all possible cutoff values. The blue line shows the specificity. The red line shows the sensitivity. In A, I show the cutoff values for C-reactive protein. In B, I show the cutoff values for procalcitonin. In C, I show the cutoff values for lytA. The ideal plot would see the blue line quickly approach one with relatively little decline in red. Unfortunately, that behavior is not present for these tests.

### Standard approach to identify a cutoff value for *S. pneumoniae*

For the three statistically significant biomarkers, I found that each filled a similar receiver operator characteristic (ROC) tract (Figs. 3A, 3B, and 3C). Each biomarker diverged far from the diagonal and displayed a similar tradeoff between true positives and false positives. As expected, it was difficult to find the ideal dichotomous cutoff by ROC alone. Therefore, I computed the Youden index and plotted it both against the traditional false positive rate and against the more useful biomarker concentration (Fig. 3). According to the Youden index, the optimal false positive rate for CRP was 0.3 and the optimal biomarker cutoff was 16.67 *mg/dL.* For procalcitonin, the optimal dichotomous false positive rate was 0.34 and the optimal biomarker cutoff was 2.22 *ng/mL.* For lytA, the optimal false positive cutoff was 0.27 and the optimal concentration was 4.47 *log_10_ copies/mL* or 2.95× 10^4^ *copies/mL.* Each cutoff was significantly higher than that used to diagnosis anything in the HIV-uninfected population. Therefore, HIV patients were either hyper-responders for inflammatory biomarkers or, more likely, *S. pneumoniae* caused a much larger inflammatory reaction than the average for all bacterial infections.

**Figure 3:**
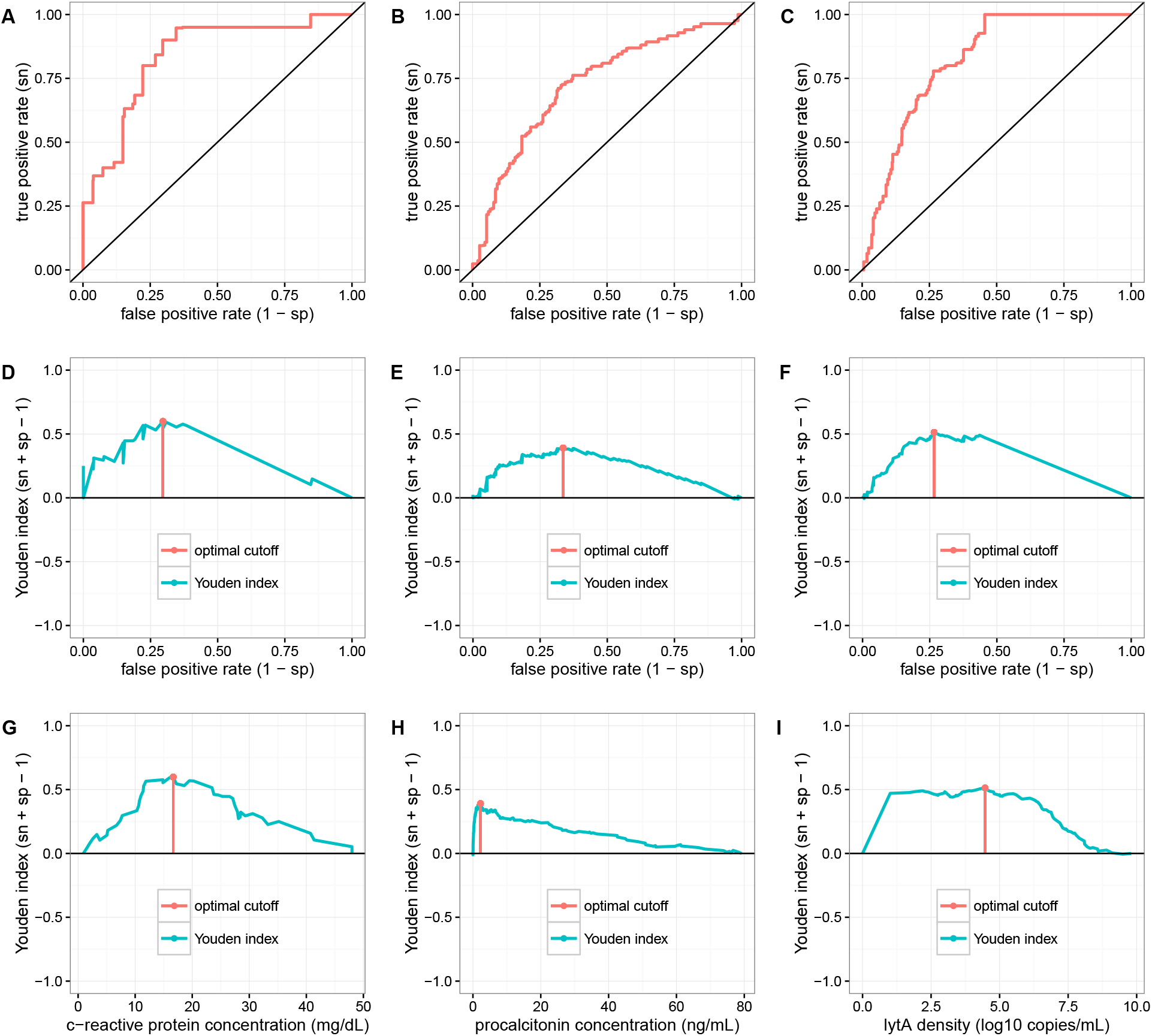
Receiver Operating Characteristic (ROC) plots and Youden index plots of each biomarker. In A, I show the ROC curve for C-reactive protein. In B, I show the ROC curve for procalcitonin. In C, I show the ROC curve for lytA. In D, I show the Youden index versus false positive rate for C-reactive protein. In E, I show the Youden index versus false positive rate for procalcitonin. In F, I show the Youden index versus false positive rate for lytA. In G, I show the Youden index versus the actual test values for C-reactive protein. In H, I show the Youden index versus the actual test values for procalcitonin. In I, I show the Youden index versus the actual test values for lytA. The Youden index is a measure of how informative each biomarker cutoff is for pneumococcal etiology. In red overlaying blue, I show the optimal dichotomous cutoff for each biomarker. The optimal value for C-reactive protein is 16.67 *mg/dL*, for procalcitonin is 2.22 *ng/mL*, and for lytA is 4.47 *log_10_ copies/mL* or 2.95 × 10^4^ *copies/mL.* By Youden index, any value of lytA less than 5 *log_10_ copies/mL* or 10^5^ *copies/mL* is essentially identical.

### Bayesian approach to identify the optimal cutoff value

In contrast to the standard approach that focuses on applying various statistical tests with a probability of incorrectness, a Bayesian approach couches the problem itself in terms of probabilities. Thus, every question to be answered requires a prior probability. In clinically-oriented statistics, the prior probability is often the disease prevalence in the naïve case or the pretest probability when more information is available. With a pretest probability, Bayesian statistics updates the probability of an event as new evidence becomes available. In clinical statistics, the easiest method to update the probability is via likelihood ratios to update pretest odds. In the case of dichotomous cutoffs, there is always a likelihood ratio positive (*LR*+) to update the odds after a positive test and a likelihood ratio negative (*LR–*) to update the odds after a negative test.

I calculated the likelihood ratio positive and likelihood ratio negative as well as the 95% confidence interval for each biomarker (Fig. 4). The three biomarkers all had an extensive range where the confidence band did not overlap one, which implies statistical significance. The lytA biomarker displayed the longest and most valuable range of likelihood ratio negative values; thus, a negative lytA test at most cutoffs lowered the posttest probability of pneumococcal infection. By contrast, there were few cutoff values for procalcitonin where a negative test lowered the posttest probability of pneumococcal pneumonia. Also important for clinical use, there were significant upper boundary effects for each of these tests. Although it may be unintuitive, both PCT and lytA seemed to have a threshold above which positive tests on higher cutoffs did not raise the posttest probability as much as a positive test at a lower cutoff. Perhaps the sampling was not sufficient to denote the real population distribution. Alternatively, it may be the case that other pneumonia etiologies become more likely for higher cutoffs.

**Figure 4:**
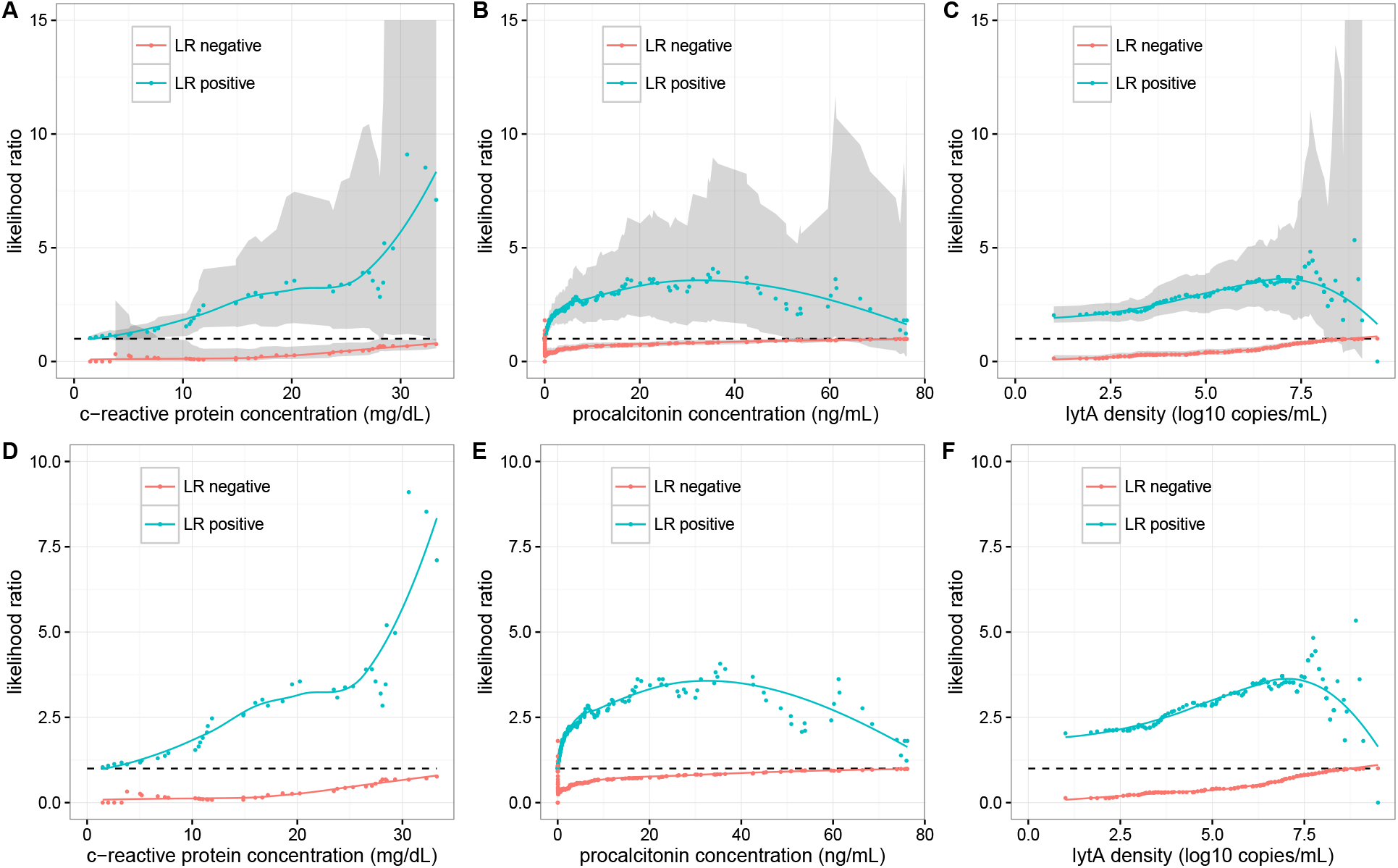
Likelihood ratios for each available cutoff in the data. The blue points show the likelihood ratio for a positive test result. The red points show the likelihood ratio for a negative test. Likelihood ratio positive values between two and five have a small effect, those between five and ten have moderate effect, and those greater than ten have large effect. The inverse is true for the likelihood ratio negative (e.g. 1/2, 1/5, etc). A likelihood ratio of one means the test is uninformative. In A, I show the values for C-reactive protein with the 95% CI in gray. In B, I show the values for procalcitonin with the 95% CI in gray. In C, I show the values for lytA with the 95% CI in gray. In D, I show the values for C-reactive protein zoomed in and removing the 95% CI. In E, I show the values for procalcitonin zoomed in and removing the 95% CI. In F, I show the values for lytA zoomed in and removing the 95% CI. In black dash, I show the *y* = 1 line. The limited sample size for C-reactive protein make it difficult to assess the upper limit. The traditional cutoff for *procalcitonin* = 0.5 *ng/mL* may need to be adjusted upward to improve clinical yield. The cutoff of *lytA* = 8 × 10^3^ *copies/mL* may be lower than ideal; the primary value of lytA compared to procalcitonin is in lytA’s negative predictive value. For PCT, *LR*— would be maximized nearer the lower detectable limit with little loss in positive predictive value.

Rather than the Youden J index, Bayesian statistics suggests a very different manner of identifying the optimal dichotomous cutoff. From a Bayesian perspective, the value of a particular cutoff on a clinical test is most directly understood as the maximum difference between the posttest probability after a postive test and the posttest probability after a negative test. Thus, the most valuable test is the one that most dramatically moves the posttest probability.

I used the pretest probability in the sample along with the *LR*+ and *LR*– to plot the predicted posttest probabilities for every possible cutoff in the data (Fig. 5). If the cutoff were set to a point on the *x*-axis, any positive test for that cutoff had the plotted posttest positive probability and any negative test had the plotted posttest negative probability. Then, to find the optimal cutoff I subtracted the posttest negative probability from the posttest positive probability. In dramatic contrast to the Youden index, this calculation showed that the vast majority of available cutoffs were essentially identical. For CRP, although the optimal value was 30.6 *mg/dL*, any value greater than 10 *mg/dL* produced a similar posttest probability difference. Likewise for PCT, the optimal cutoff was 17.4 *ng/mL*, but cutoffs up to 40 *ng/mL* were similarly informative. Interestingly, the optimal cutoff for lytA was exactly the same value as that found by the Youden index, 4.47 *log*_10_ *copies/mL* or 2.95 × 10^4^ *copies/mL.*

**Figure 5:**
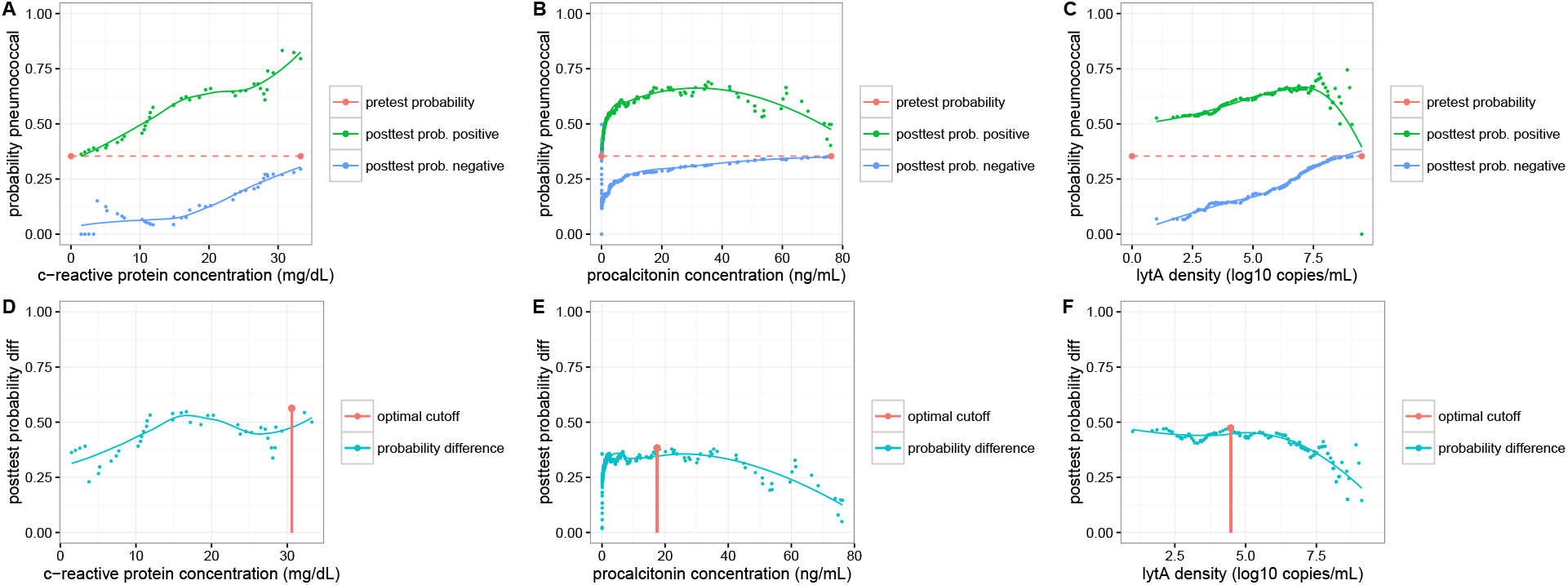
Probability plots for each biomarker. In A, I show the posttest probability calculations for C-reactive protein. In B, I show the posttest probability calculations for procalcitonin. In C, I show the posttest probability calculations for lytA. In D, I show the difference between the positive posttest probability and negative posttest probability for C-reactive protein. In E, I show the difference between the positive posttest probability and negative posttest probability for procalcitonin. In F, I show the difference between the positive posttest probability and negative posttest probability for lytA. In red dash, I show the pretest probability. In terms of Bayesian information, the ideal dichotomous cutoff is that value that maximizes the difference between the positive posttest probability and the negative pottest probability. Thus, the optimal cutoff for C-reactive protein is 30.6 *mg/dL*, for procalcitonin is 17.4 *ng/mL*, and for lytA is 4.47 *log_10_ copies/mL* or 2.95 × 10^4^ *copies/mL.* The optimal cutoff of lytA by Bayesian yield is almost identical to that by Youden index. However, as with Youden index, any value less than 5 *log_10_ copies/mL* is essentially identical. In addition, by Bayesian yield there is no clear ideal value for any biomarker; many values perform similarly well.

### Quantitative likelihood ratios improve probability calculations

Although Bayesian analysis suggests somewhat different cutoffs, its real value lies in the ability to easily incorporate quantitative data into probabilistic belief. Thus, I computed a continuous likelihood ratio for each biomarker (Fig. 6). I found that PCT displayed very different behavior from that of either CRP or lytA. There were few values of PCT that lowered the pretest probability of pneumococcal etiology. By contrast, depending on the result CRP and lytA could either raise or lower the posttest probability of infection. Unfortunately, in every case computing continuous likelihood ratios smoothed over potentially informative edge effects in the data. Therefore, I calculated the posttest probabilities using both the combined and the continous likelihood ratio values (Fig. 7). Throughout most biomarker values, continuous and combined likelihood ratios produced remarkably similar posttest probabilities. For CRP and lytA, the posttest probability calculated with the combined likelihood ratio was essentially identical to that calculated with the continuous logistic function; the only significant deviations were near the ends of the test range. For PCT, there were more significant differences between the combined and continuous posttest probabilities. Using the combined likelihood ratios, procalcitonin levels greater than 20 *ng/mL* did not produce higher posttest probabilities. Thus, it appeared that higher values of CRP and lytA encoded ever increasing clinical probabilities while PCT displayed a more narrow informative range.

**Figure 6:**
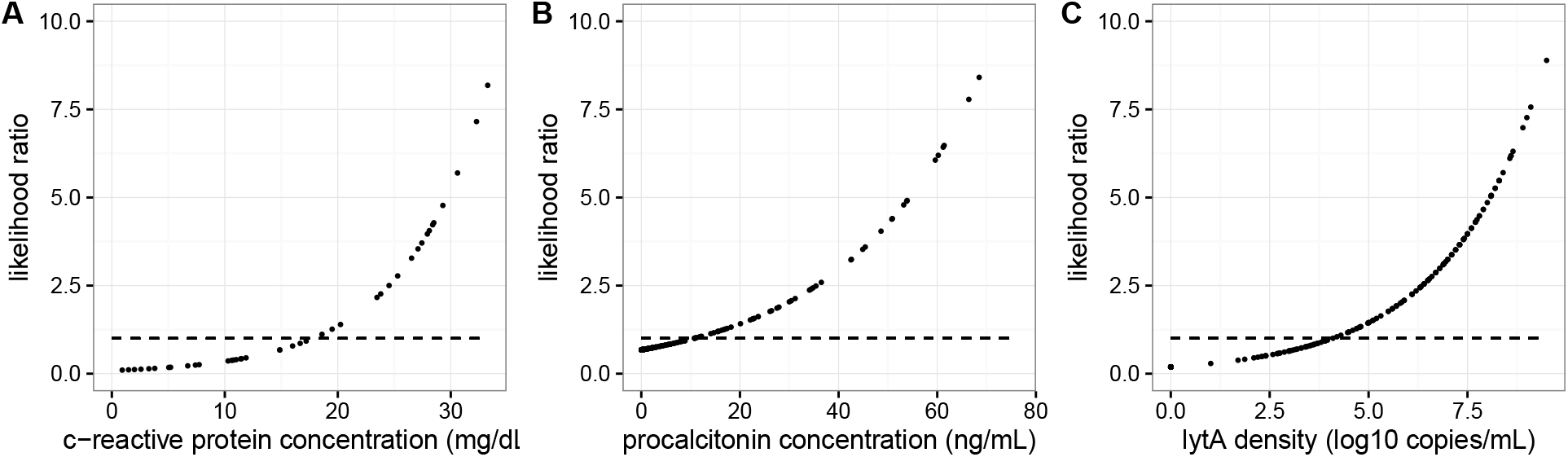
Continuous likelihood ratios for several pretest probabilities [26]. In A, I show the likelihood ratios for C-reactive protein with the actual pretest probability in the data. In B, I show the likelihood ratios for procalcitonin with the actual pretest probability in the data. In C, I show the likelihood ratios for lytA with the actual pretest probability in the data. The dashed line is the uninformative likelihood ratio at *LR* = 1.

**Figure 7:**
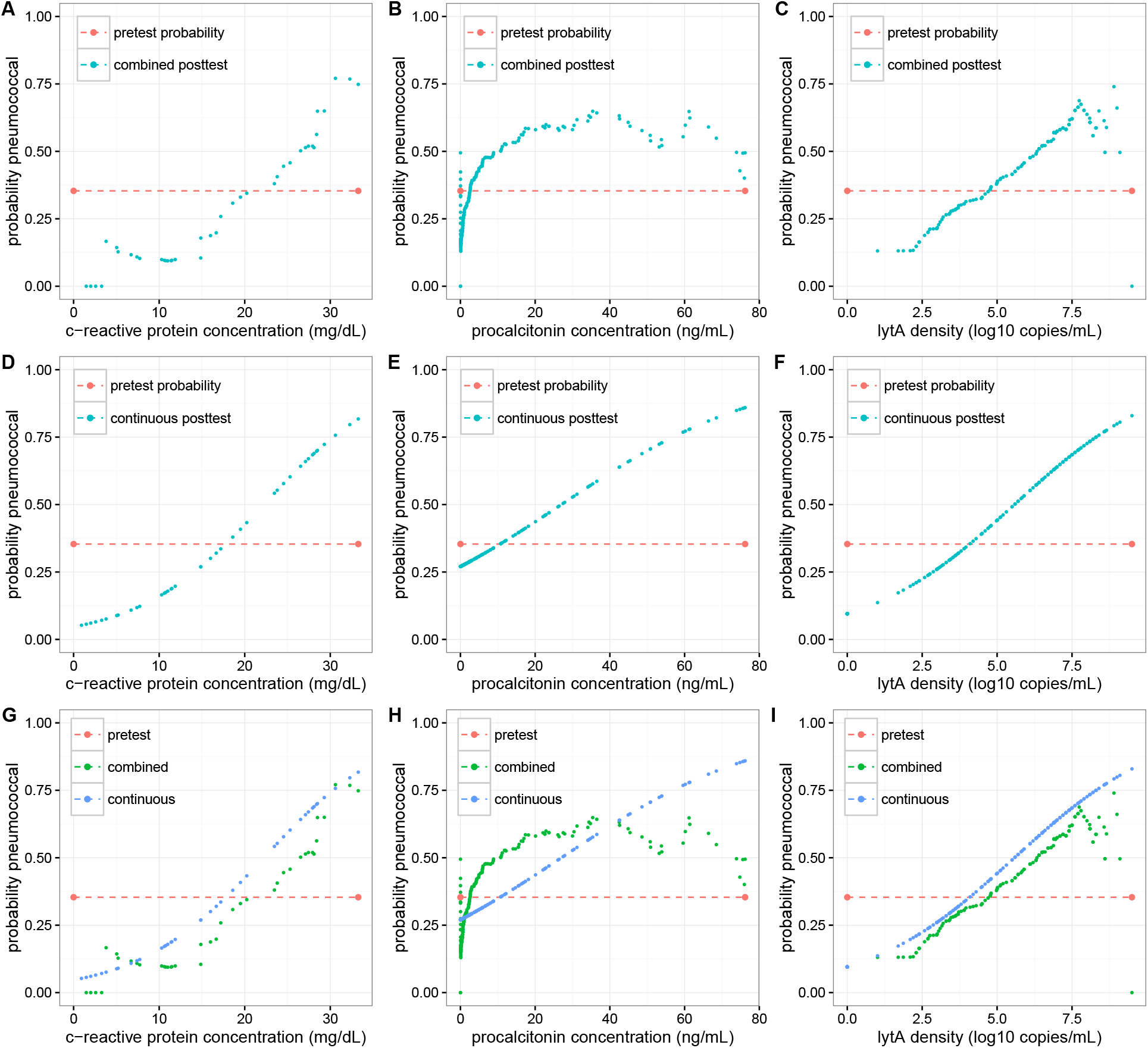
Posttest probability calculations using combined likelihood scores. In A, I show the calculated posttest probability combining the LR+ and *LR—* values at each point for C-reactive protein. In B, I show the calculated posttest probability combining the LR+ and *LR—* values at each point for procalcitonin. In C, I show the calculated posttest probability combining the LR+ and *LR—* values at each point for lytA. In D, I show the calculated posttest probability using the continuous *LR* values for C-reactive protein. In E, I show the calculated posttest probability using the continuous *LR* values for procalcitonin. In F, I show the calculated posttest probability using the continuous *LR* values for lytA. In G, I show A and D overlay to display their correlation. In H, I show B and E overlay to show their correlation. In I, I overlay C and F to show their correlation. The bottom row shows that one can empirically calculate posttest probability without the need for logistic regression. In red dash, I show the pretest probability of pneumococcal infection.

In terms of clinical utility, my analysis showed that CRP and lytA were broadly capable of making a pneumococcal diagnosis either highly likely or highly unlikely. A CRP test result of less than 5 *mg/dL* meant the posttest probability of pneumococcal etiology was less than 15%. That was also the case for lytA values of less than 2.5 *log*_10_ *copies/mL*. On the other end of the spectrum, relatively high levels of CRP and lytA resulted in posttest probabilities approaching 70%. Also, there was a clear warning to those setting dichotomous cutoffs on these tests. For each biomarker, the combined posttest probability intersected the pretest probability at almost exactly the optimum value suggested by the Youden index. If the cutoff was set at that value, test results nearby were almost completely uninformative (manipulating the available web application may show this most clearly). Since many or most test results fall in a relatively narrow range near the optimal cutoff, setting any dichotomous cutoff could be extremely misleading for clinicians interpreting the test results.

## Discussion

In this study, I showed that each of the three biomarkers CRP, PCT, and lytA displayed a moderate connection to pneumonia etiology. Of the three, CRP and lytA were similarly valuable in determining pneumococcal etiology whereas PCT displayed the least diagnostic power. I showed that existing cut points for all three tests were not the ideal cutoffs for diagnosing pneumococcal pneumonia in HIV patients. Thus, cutoffs need to be adjusted for etiology discrimination. Then, I utilized a Bayesian framework to calculate the likelihood ratios for every possible cutoff. Throughout each clinical test range, the *LR*+ and *LR*– were statistically significant. Next, I showed that the standard approach to cut point identification failed to find the clinical cutoff that optimized for posttest probability yield for CRP and PCT. Furthermore, for lytA I found that the Youden index and the optimal Bayesian cutoff were identical, and yet different from the existing cut point. Then, using two different methods, I showed that it was unnecessary and wasteful to discard the quantitative data inherent in these tests. For each biomarker, there was no indication of a cut point that lead to categorical diagnoses; lower values led to a lower posttest probability while higher values led to a higher posttest probability. I showed that even in the absence of a logistic regression model one can easily calculate an empirical combined likelihood ratio for every possible test result. Finally, I showed that the quantitative value of these tests can be dramatic. For CRP and lytA, the posttest probability of pneumococcal pneumonia is greater than 65% with high values of each test. By contrast, when test values approach zero, the posttest probability of pneumococcal pneumonia vanishes to less than 15%. Thus, with a Bayesian approach these biomarkers can dramatically lower the uncertainty of *S. pneumoniae* infection following chest x-ray confirmed pneumonia in HIV patients.

Although there are a number of prior studies that evaluate the value of various biomarkers in the diagnosis and treatment of pneumonia, relatively few use biomarkers to help diagnose a particular infectious etiology. Furthermore, almost none give concrete guidance to clinicians regarding the posttest probability of a particular organism. In most cases, investigators note a “statistically significant” difference in the distribution of test results between healthy and diseased populations. For example, many papers suggest CRP [8,11] and PCT [11,13–17] can add value to either the diagnosis or treatment of a wide range of ailments. Specifically, PCT can shorten the duration of clinical treatment and improve time-to-treatment. I found only one paper where PCT proved capable of distinguishing between typical and atypical pneumonia [13]. By contrast, there are several studies showing that lytA can differentiate pneumococcal from non-pneumococcal pneumonia [22]. Unfortunately, beyond suggesting a single dichotomous cutoff, none of those studies provide information regarding the probability of *S. pneumoniae* infection. Therefore, they fail to answer the most relevant clinical question... with this test result, how much more or less likely is *S. pneumoniae* infection? Even in studies where the relative risk is calculated, the inability to incorporate a pretest probability into the relative risk ratio means that the accuracy of posttest estimation is severely diminished. Hopefully my analysis and/or open source web application can provide direct guidance to clinicians treating HIV patients.

In addition to the interpretation challenges with clinical biomarkers, there may be further issues with the standard approach to calculating clinical cutoffs. To find a cutoff, typically investigators use some permutation of the ROC curve. Briefly, ROC curves plot the true positive rate (i.e. *sensitivity)* against the false positive rate (i.e. 1 — *specificity).* The curve provides the clinical cost-benefit ratio of the loss in test specificity to gain in test sensitivity. Intuitively, a curve far from the diagonal indicates a good test and one near the diagonal indicates a poor test. Formally, there are two common ways to compute the optimal dichotomous cutoff. One, the area under the curve (AUC) approach, is intuitively simple; it finds the biomarker value that places the curve closest to the (0,1) point. Thus, AUC maximizes the true positive rate and simultaneously minimizes the false positive rate. However, work in basic statistics shows that the AUC calculation method requires the use of an additional term that causes the statistic to deviate from the ideal unweighted classification/misclassification point [24]. By contrast, the Youden J index (i.e. *max[y(x)* – *x*] on the ROC) always provides the optimal dichotomous cutoff [24]. Although the two quantities agree in many situations, the Youden index is the preferred metric when they do not. When the goal of a study is to find the ideal cutoff, there is no other weighting (e.g. economic weighting) involved, and the data exclude the possibility of Bayesian analysis, I recommend using the Youden J index rather than the more common AUC method.

Although dichotomous cutoffs for clinical tests are the tradition, there is no reason to prefer a single cutoff. To the contrary, many clinical tests provide quantitative data and in most cases setting a single cutoff on a continuous variable throws out useful clinical information. There are several available approaches to retain at least some quantitative information. One method involves calculating so called multilevel likelihood ratios. In that case, the investigator defines more than a single cutoff to produce several ordinal categories [28–30]. Such an approach is most useful when the data itself is ordinal (e.g. a pain scale from 1 to 10 or a Likert scale where the numbers can be ranked but the distance between adjacent numbers has no meaning). However, most modern clinical tests contain either interval (e.g. temperature in fahrenheit where zero degrees has no physical meaning) or ratio (e.g. weight, height, etc. where the number zero has a clear physical meaning) data types. In such cases, investigators can calculate a likelihood ratio for each possible test result. Although there is at least one existing approach to make such a computation, it requires first fitting a logistic regression curve and subsequently using coefficients of the fit. Although that approach may be relatively simple for biostatistics professionals, it is far from obvious how it may be applied more broadly. More importantly, the logistic function itself makes certain fundamental assumptions about the relationship between predictor and response variables. For example, it assumes that there are just two response states; an assumption that essentially all biomarkers violate. For most clinical tests, there are a number of possible etiologies that might fit into various portions of the test range. For example, in this study the combined PCT curve above 10 *ng/mL* deviates significantly from that calculated with logistic regression. Moreover, the combined likelihood ratio actually declines past 30 *ng/mL.* One possible and likely interpretation of this result is that other, more inflammatory infections like *H.influenzae* or *S.aureus* dominate higher ranges of the PCT test. By contrast, lytA being very specific for *S.pneumoniae* shows no such deviation until the edge of the test range where low sampling distorts the otherwise strong correlation. Thus, there are clear practical reasons to prefer the combined likelihood ratio calculation. Whatever the case, to improve the usability of clinical studies, there is no reason to throw out clinical information in favor of simple cutoffs. Continuous or combined likelihood ratios provide the ideal explanatory framework for reporting new clinical tests.

## Acknowledgments

I would like to acknowledge the epidemiologists and experimentalists who collected this data. I would like to highly commend them for publicly releasing the data. If every scientist were so forthcoming we could witness a new revolution in the biosciences.

